# Planarian LDB and SSDP proteins scaffold transcriptional complexes for regeneration and patterning

**DOI:** 10.1101/2023.02.07.527523

**Authors:** Taylor Medlock-Lanier, Kendall B. Clay, Rachel H. Roberts-Galbraith

**Author notes:** Authors contributed equally (Taylor Medlock-Lanier, Kendall B. Clay).

## Abstract

Sequence-specific transcription factors often function as components of large regulatory complexes. LIM-domain binding protein (LDB) and single-stranded DNA-binding protein (SSDP) function as core scaffolds of transcriptional complexes in animals and plants. Little is known about potential partners and functions for LDB/SSDP complexes in the context of tissue regeneration. In this work, we find that planarian LDB1 and SSDP2 promote tissue regeneration, with a particular function in mediolateral polarity reestablishment. We find that LDB1 and SSDP2 interact with one another and with characterized planarian LIM-HD proteins Arrowhead, Islet1, and Lhx1/5-1. *SSDP2* and *LDB1* also function with *islet1* in polarity reestablishment and with *lhx1/5-1* in serotonergic neuron maturation. Finally, we show new roles for LDB1 and SSDP2 in regulating gene expression in the planarian intestine and parenchyma; these functions may be LIM-HD-independent. Together, our work provides insight into LDB/SSDP complexes in a highly regenerative organism. Further, our work provides a strong starting point for identifying and characterizing potential binding partners of LDB1 and SSDP2 and for exploring roles for these proteins in diverse aspects of planarian physiology.

## Introduction

Dynamic gene expression drives consequential changes in cellular behavior during development and regeneration. Though DNA-binding transcription factors are sometimes considered in isolation, most such factors function *in vivo* in modular transcriptional complexes that bring together numerous gene regulatory proteins to elicit changes in chromatin architecture and transcriptional activity. LIM-domain binding protein (LDB)—also known as CLIM, NLI, and CHIP—scaffolds transcriptional complexes with the help of Single-stranded DNA-binding protein (SSDP) (Chen *et al.*, 2002; van Meyel *et al.*, 2003). LDB and SSDP scaffolded complexes are ancient, present in diverse animals, and are related to similar scaffolding proteins (SEUSS/LEUNIG) that work together to organize gene regulatory complexes in plants (Franks *et al.*, 2002; van Meyel *et al.*, 2003; Sridhar *et al.*, 2004).

As core transcriptional regulatory factors in gene expression across broad cell types, LDB and SSDP orthologs play diverse roles in development, particularly in formation of the head and brain (Matthews and Visvader, 2003). *Drosophila* Chip was identified in a genetic screen as a regulator of wing morphology (Morcillo *et al.*, 1996; Morcillo *et al.*, 1997), and *Drosophila* SSDP is essential, with mutants lacking maternal and zygotic SSDP dying during larval development (Chen *et al.*, 2002). *C. elegans SAM-10* (ortholog of SSDP) and *ldb-1* promote normal gene expression in neurons during maturation, modulating synapse and neurite formation (Cassata *et al.*, 2000; Zheng *et al.*, 2011). Zebrafish SSDP and CLIM/LDB paralogs function in neural patterning, eye formation, and sensory axon formation (Becker *et al.*, 2002; Zhong *et al.*, 2011). Mice lacking *ssdp1* develop with aberrant head morphogenesis; the initial mouse mutant in *ssdp1* was named *headshrinker* (Nishioka *et al.*, 2005). A proline-rich region within SSDP1 was also found to be essential for proper head development (Enkhmandakh *et al.*, 2006). Mice lacking LDB1 also develop with severe anterior truncations and defects in head development, as well as other pleiotropic phenotypes including heart defects and patterning problems (Mukhopadhyay *et al.*, 2003), while murine LDB2 plays overlapping *and* unique roles in neural development (Gueta *et al.*, 2016; Leone *et al.*, 2017). Finally, LDB proteins can function in chromatin looping and recruitment of complexes to modify chromatin (Krivega *et al.*, 2014; Caputo *et al.*, 2015; Lee *et al.*, 2016; Magli *et al.*, 2019).

In animals, LDB/SSDP complexes often include LIM-homeodomain proteins and other sequence-specific DNA-binding transcription factors that direct the complex to specific regions of the genome (Matthews and Visvader, 2003) (Fig. 1A). LDB itself can function as a dimer or trimer, resulting in the potential bridging of two transcription factors together to form homo- or hetero-multimeric DNA-binding structures (Jurata *et al.*, 1998; Cross *et al.*, 2010; Wang *et al.*, 2020). Though perhaps best known for binding LIM-HD proteins (Agulnick *et al.*, 1996; Morcillo *et al.*, 1997; Hobert and Westphal, 2000; van Meyel *et al.*, 2003; Yasuoka and Taira, 2021), LDB/SSDP complexes can also include transcriptional regulators that do *not* possess LIM domains. Additional LDB/SSDP-associated transcription factor proteins have been thoroughly documented in *Drosophila*, where they include Pygopus, Bicoid, Pannier, Groucho, and Ftz (Torigoi *et al.*, 2000; Bronstein *et al.*, 2010; Fiedler *et al.*, 2015). Cooperation between LDB, SSDP, and transcription factors leads to varied outcomes. For example, murine Lim1 plays a critical role in head organization in development (Shawlot and Behringer, 1995), similar to functions of mouse LDB1 and SSDP. And murine Lhx2 cooperates with LDB1 for a narrower role during several neurodevelopmental processes, including regionalization of the hippocampus, development of olfactory neurons, and regulation of neurogenic vs. gliogenic choices in the retina (de Melo *et al.*, 2018; Monahan *et al.*, 2019; Kinare *et al.*, 2020).

**Figure 1.**
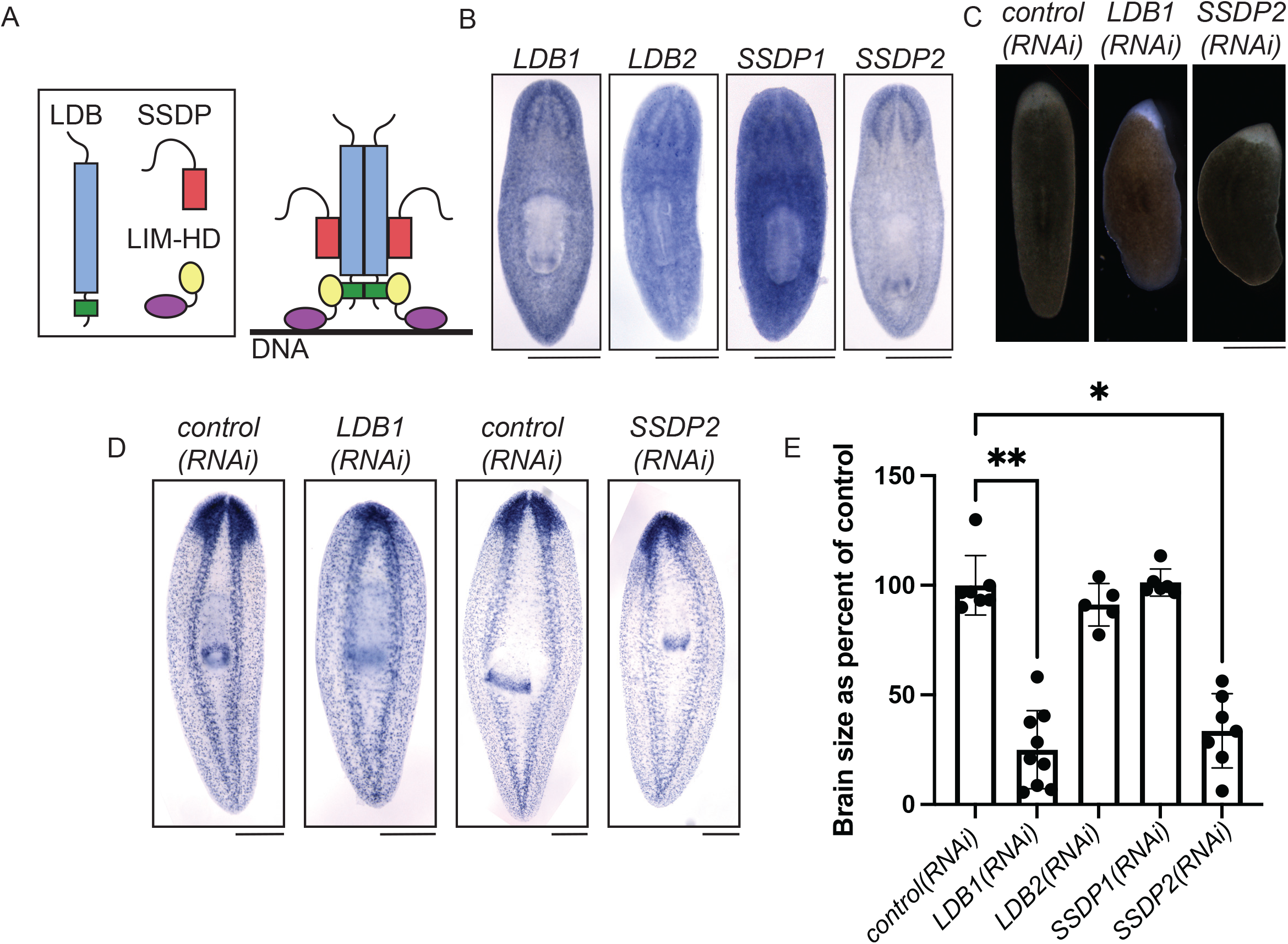
1) Planarian SSDP and LDB homologs promote regeneration. A) Diagram showing domain architecture in canonical LDB transcriptional complexes (Enkhmandakh *et al.*, 2006; Bronstein *et al.*, 2010). LDB=LIM Domain Binding protein; SSDP=Single-Stranded DNA Binding Protein; and LIM-HD=LIM-Homeodomain, where LIM is a region of homology first identified between Lin-11, Isl-1 and Mec-3. LDB proteins have a long dimerization domain (blue) and a LIM-binding domain (green). SSDP proteins have a conserved domain (red) and a long, unstructured region. LIM-HD proteins have a pair of LIM domains (yellow) and a homeodomain (purple). Complexes form to bind DNA and regulate gene expression. B) *in situ* hybridization showing expression patterns of planarian *LDB1*, *LDB2*, *SSDP1*, and *SSDP2*. All four genes are broadly expressed with some enrichment, particularly for *LDB1* and *SSDP2*, in the brain. C) Images of live planarians after RNAi, head amputation, and 6 days of regeneration. *control(RNAi)* animals have a rounded blastema with two eyespots visible (n=10). *LDB1(RNAi)* and *SSDP2(RNAi)* regenerates have pointed blastemas with zero or one eyespot (6/6 and 9/10, respectively). D). *in situ* hybridization to mark the broadly expressed transcript *ChAT* (*choline acetyltransferase;* (Nishimura *et al.*,2010)) in animals after RNAi, head amputation, and 6 days of regeneration. *LDB1(RNAi)* and *SSDP2(RNAi)* animals have smaller brains which are fused at the midline. E) Quantification of animals from (D) showing significantly smaller brains. (n=5-9; Kruskal-Wallis with Dunn’s correction for multiple comparisons.) P is an adjusted value, * indicates P≤0.05 and ** indicates P≤0.005. Scale=500 μm.

SSDP and LDB homologs regulate head organization and neural development during embryogenesis, but few studies have explored potential roles for these core regulators during regeneration. We opted to explore potential functions of LDB, SSDP, and associated transcription factors in regeneration using planarian flatworms of the species *Schmidtea mediterranea.* Planarians complete whole-body regeneration from tiny fragments of starting tissue using adult, pluripotent stem cells (for review, see (Ivankovic *et al.*, 2019)). During regeneration, planarians reestablish diverse cell types in organs that include muscle, a brain, protonephridia, and a muscular feeding organ called the pharynx (Roberts-Galbraith and Newmark, 2015). Due to their unique biology, planarians are outstanding model organisms with which to discover genetic mechanisms that regulate regeneration and stem cell biology. Planarian SSDP and LDB homologs have not been characterized, but a handful of LIM-homeodomain proteins have been studied. Planarian *islet1* plays roles in reestablishment of anterior, posterior, and midline polarity (Hayashi *et al.*, 2011; Marz *et al.*, 2013). *lhx1/5-1* directs maturation of serotonergic neurons in the central and peripheral nervous system of the planarian (Currie and Pearson, 2013). And an *arrowhead* homolog is required for reestablishment of medial brain structures, including the anterior commissure, through creation of neurons that serve as guidepost cells (Roberts-Galbraith *et al.*, 2016; Scimone *et al.*,2020).

Here, we identify and characterize roles for planarian LDB and SSDP homologs in the context of regeneration. We find that LDB1 and SSDP2 are required for proper regeneration, playing important roles in reestablishment of polarity. Using yeast two-hybrid assays, we show that LDB1 interacts with itself, with SSDP2, and with LIM-HD proteins Islet1, Lhx1/5-1, and Arrowhead. We further show that LDB1 and SSDP2 likely cooperate with binding partners Islet1 and Lhx1/5-1 in polarity reestablishment and serotonergic maturation, respectively. Broadening our study, we describe the complement of genes that encode LIM-homeodomain and LIM-only domain proteins in planarians. Finally, to begin the process of identifying potential targets of SSDP2-containing complexes, we performed RNAi and RNA-sequencing and identified dozens of genes that depend on SSDP2 for proper expression. We conclude that planarian SSDP2 and LDB1 play key roles in regulating gene expression during planarian regeneration both by cooperating with known LIM-HD partners and likely other transcriptional regulators. Our work provides an important foundation for exploring the shared and unique roles of conserved LDB/SSDP complexes in the context of robust tissue regeneration.

## Results and Discussion

### Planarian LDB1 and SSDP2 are required for blastema size and organization

To identify planarian LDB and SSDP homologs, we searched available transcriptomes (Brandl *et al.*, 2016; Rozanski *et al.*, 2019) and found two homologs each of *LDB* and *SSDP* (Supp. Tables 1, 2). We examined the expression patterns of planarian homologs and determined that *LDB1*, *LDB2*, *SSDP1*, and *SSDP2* have nearly ubiquitous expression with some potential enrichment in the planarian brain (Fig. 1B). Wide-ranging expression was also indicated in published single-cell sequencing atlases, with some specific enrichment of *LDB1* in subsets of *cathepsin^+^* cell types (Supp. Tables 1, 2, (Fincher *et al.*, 2018; Plass *et al.*, 2018)). We also note that both *LDB1* and *SSDP2* were significantly upregulated at 36 hours and 72 hours post-amputation in our prior gene expression analysis (Roberts-Galbraith *et al.*, 2016).

Next, we investigated whether planarian LDB and SSDP homologs play roles in regeneration. We knocked down *LDB1*, *LDB2*, *SSDP1*, or *SSDP2*, amputated prepharyngeally, and allowed heads to regenerate for 6 days. We did not note any regeneration defects after *LDB2(RNAi)* or *SSDP1(RNAi).* However, *LDB1(RNAi)* and *SSDP2(RNAi)* animals exhibited slower movement and regenerated with pointy head blastemas and cyclopia (Fig. 1C). We also repeated RNAi of each homolog, amputated heads, and examined brain organization at 6 days post-amputation (dpa) via *in situ* hybridization (ISH) with a *choline acetyltransferase* (*ChAT*) riboprobe (Nishimura *et al.*,2010). We found that brain regeneration proceeded normally after *LDB2(RNAi)* or *SSDP1(RNAi).* However, *LDB1(RNAi)* and *SSDP2(RNAi)* animals regenerated with significantly smaller brains that were fused at the midline (Fig. 1D-E). We conclude that planarian LDB1 and SSDP2 are important for head regeneration, just as SSDP and LDB homologs are important for head development in mice (Mukhopadhyay *et al.*,2003; Nishioka *et al.*, 2005). We opted to focus on LDB1 and SSDP2 for the remainder of our studies.

### Smed-LDB1 interacts with SSDP2 and LIM-HD binding partners

In other animals, LDB homologs scaffold protein complexes and bind to SSDP and LIM-HD proteins to regulate transcription (Fig. 1A, (Matthews and Visvader, 2003)). To test whether planarian LDB1 binds similarly to SSDP and LIM-HD proteins, we turned to yeast two-hybrid analysis. We decided to focus on characterized planarian LIM-HD proteins Lhx1/5-1, Islet1, and Arrowhead (Hayashi *et al.*, 2011; Currie and Pearson, 2013; Marz *et al.*, 2013; Roberts-Galbraith *et al.*, 2016; Scimone *et al.*, 2020). We cloned full-length (FL) versions of each gene into yeast two-hybrid vectors to create fusions of planarian proteins with either the DNA-binding domain (BD, Bait) or the activation domain of GAL4 (AD, Prey) (Chien *et al.*, 1991). We also cloned short (Sh) versions of each gene that included only the domains relevant for known protein-protein interactions (Fig. 2A). For LDB1, we cloned the dimerization domain and LIM-binding domain. For SSDP2, we cloned the conserved N-terminal domain. We cloned the N-termini of Islet1, Lhx1/5-1, and Arrowhead (Ah), including both LIM domains for each LIM-HD protein.

**Figure 2.**
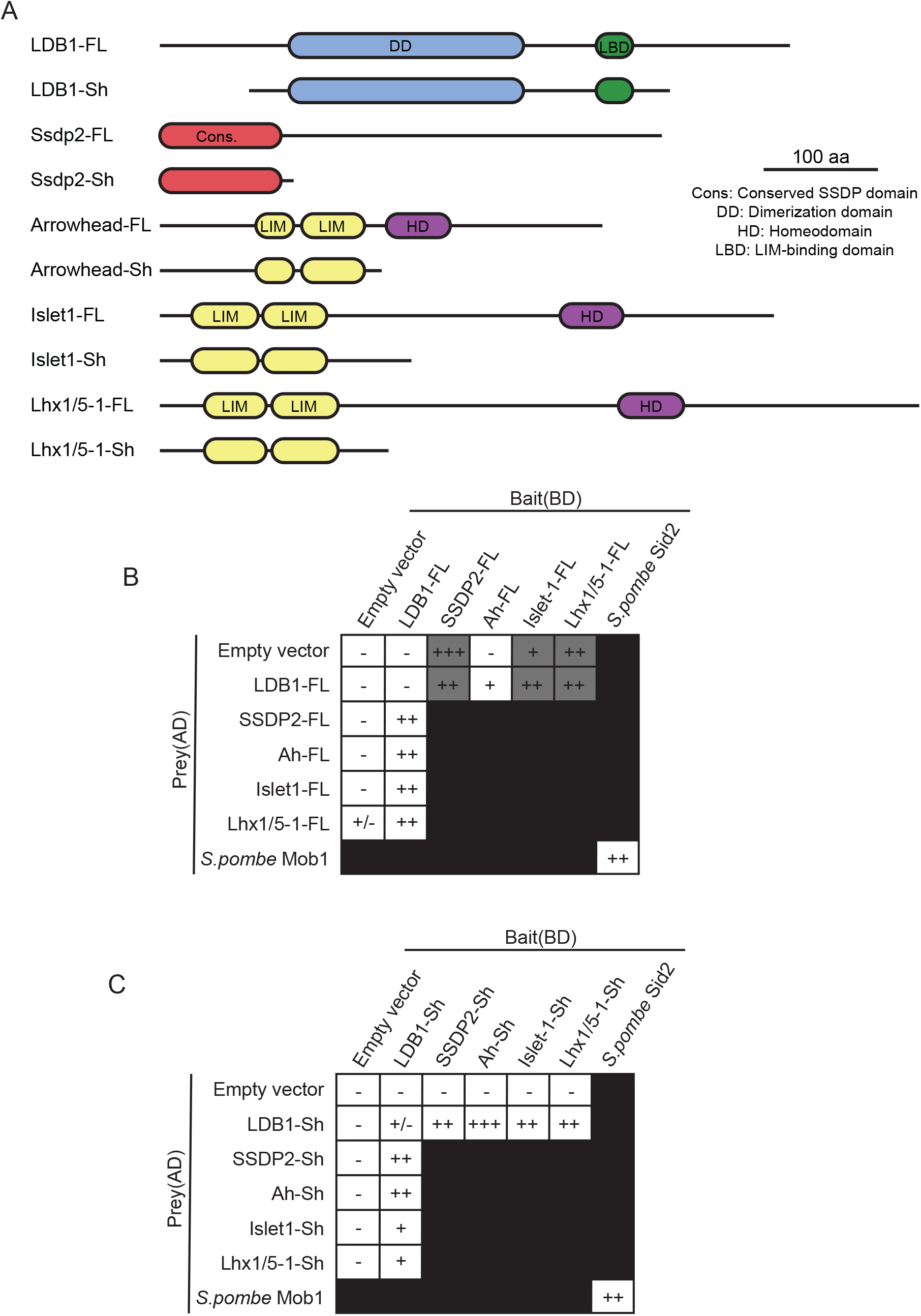
2) Planarian LDB1 interacts with SSDP2, as well as the LIM domains of Arrowhead, Islet, and Lhx1/5-1. A) Diagram showing domain architecture for LDB1, SSDP2, Arrowhead, Islet, and Lhx1/5-1. “FL” indicates full length protein and “Sh” indicates a shorter truncation used in subsequent analyses. The color scheme used is the same as in Fig. 1A. B) Yeast two-hybrid results showing the interaction of full-length LDB1 with full-length SSDP2, Arrowhead, Islet, and Lhx1/5-1. Full-length Ssdp2, Islet, and Lhx1/5-1 exhibited transactivation when used as the Bait (dark grey), so these pairings were not used for the final assessment. Full length planarian LDB1 did not show interaction with itself, despite known homodimerization in other models. (- indicates no interaction; +/- indicates weak growth; +, ++, or +++ indicates growth on restrictive medium). C) Yeast two-hybrid results showing that the core domains of LDB1 (LDB1-Sh) interact with the conserved domain of SSDP1 and the LIM domains of Arrowhead, Islet, and Lhx1/5-1. In this experiment, we did note a weak self-interaction between the core domains of LDB1, which could support some dimerization of the protein. *S. pombe* Mob1 and Sid2 were used as controls (Balasubramanian *et al.*, 1998; Hou *et al.*, 2004). Cell growth for B and C is shown in Supp. Fig. 1A-B.

In this yeast two-hybrid system, pairs of plasmids are transformed into an auxotrophic strain of *Saccharomyces cerevisiae.* Interaction between bait and prey plasmids enables activation of gene expression to allow growth on drop-out media lacking histidine and adenine supplements. With this assay, we determined that fulllength planarian LDB1 interacted with planarian SSDP2, Arrowhead, Lhx1/5-1, and Islet1 (Fig. 2B, Supp. Fig. 1A). We did not see evidence of LDB1 self-interaction using full-length protein, contrary to studies of LDB in other organisms. We were only able to establish binding of LDB1 to partners using LDB1 as bait, due to transactivation of planarian SSDP2, Islet1, and Lhx1/5-1 when fused with the DNA-binding domain (BD, bait) (Fig. 2B, Supp. Fig. 1A). This experimental complication may indicate that fulllength SSDP2, Islet1, and Lhx1/5-1 can directly recruit *S. cerevisiae* proteins to promote transcription.

We confirmed interactions and narrowed interaction domains using our “Sh” constructs. We found that the conserved interior of planarian LDB1 interacts with the conserved N-terminus of SSDP2 (Fig. 2C, Supp. Fig. 1B). We also found that conserved LDB1 domains interact with the LIM domains of Arrowhead, Islet1, and Lhx1/5-1 (Fig. 2C, Supp. Fig. 1B). We additionally detected some interaction of the short construct of LDB1 with itself in this assay (Fig. 2C, Supp. Fig. 1B), which may indicate that planarian LDB1 does indeed self-interact. We did not observe transactivation of any short construct, which indicates that the excluded regions of SSDP2 and LIM-HD proteins are responsible for transactivation in this system.

We conclude that planarian LDB1 and SSDP2 interact similarly to their homologs in other animals (Fig. 1A). Further, LDB1 can interact with the LIM domains of planarian Islet1, Lhx1/5-1, and Arrowhead. Smed-LDB1 also self-interacts, albeit weakly in this assay. These protein-protein interactions demonstrate potentially conserved organization within planarian LDB1/SSDP2 complexes, though studies in planarian cells or extract will be required to demonstrate *in vivo* interactions and study complex composition in more detail.

### LDB1 and SSDP2 cooperate with LIM-HD proteins Islet1 and Lhx1/5-1

Based on our determination that planarian LDB1 can bind to Arrowhead, Islet1, and Lhx1/5-1, we next sought to understand whether planarian LDB1 and SSDP2 cooperate with these LIM-HD proteins. We first investigated whether planarian LDB1 and SSDP2 cooperate with Lhx1/5-1. Lhx1/5-1 is expressed in cells throughout the central and peripheral nervous system and promotes maturation of serotonergic neurons (Fig. 3A, (Currie and Pearson, 2013)). Knockdown of *lhx1/5-1* reduces expression of *tryptophan hydroxylase* (*TPH*), a marker of peripheral serotonergic neurons, without affecting *TPH^+^* cells in the eyes (Fig. 3B, inset, (Nishimura *et al.*, 2007; Currie and Pearson, 2013)). To assess whether *LDB1* or *SSDP2* is required for the function of *lhx1/5-1* in serotonergic neurons, we performed RNAi and examined *TPH* expression (Fig. 3C). We found that expression of *TPH* in the peripheral nervous system was reduced after *LDB1(RNAi)* or *SSDP2(RNAi)* (Fig. 3C, insets). We also detected cyclopia using the *TPH* marker that was not present after *lhx1/5-1(RNAi)* but was present after *islet1(RNAi)*, as has been previously reported (Fig. 3B-C, (Marz *et al.*,2013)). We conclude that LDB1 and SSDP2 overlap functionally with Lhx1/5-1 in promoting gene expression in peripheral serotonergic neurons, supporting the idea that these proteins work in a transcriptional complex in this process.

**Figure 3.**
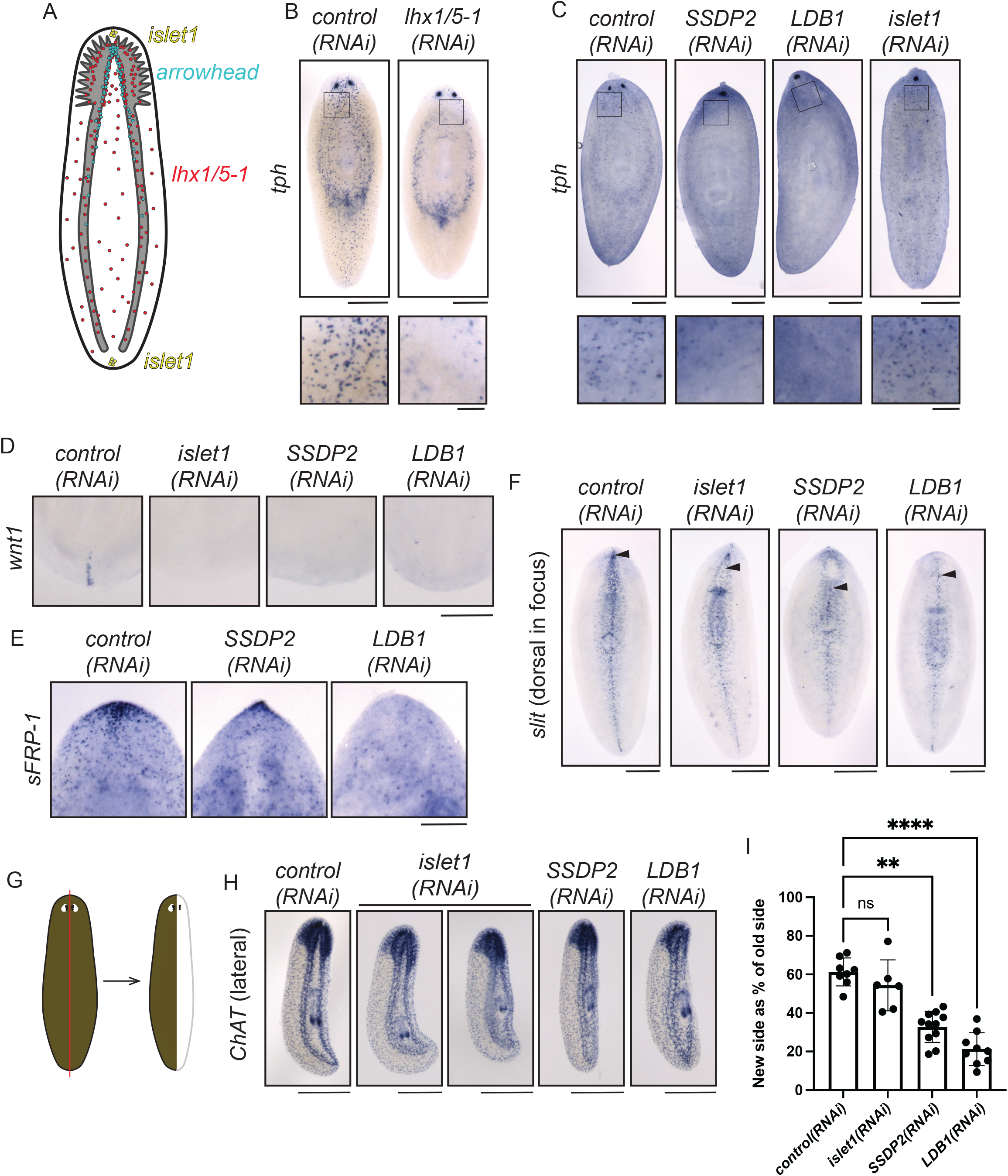
3) *SSDP2(RNAi)* and *LDB1(RNAi)* phenotypes indicate functional overlap with Islet1 and Lhx1/5-1. A) Diagram depicting published expression patterns of planarian LIM-HD-encoding genes *islet1* (Hayashi *et al.*, 2011; Marz *et al.*, 2013), *arrowhead* (Roberts-Galbraith *et al.*, 2016), and *lhx1/5-1* (Currie and Pearson, 2013). B) *control(RNAi)* and *lhx1/5-1(RNAi)* animals underwent *in situ* hybridization with *TPH* (*tryptophan hydroxylase;* (Nishimura *et al.*, 2007)) 6 days after head amputation. Fewer peripheral *TPH^+^* neurons are seen after *lhx1/5-1(RNAi)* (see inset at bottom), as seen in a prior study (Currie and Pearson, 2013). C) control(RNAi), *SSDP2(RNAi)*, *LDB1(RNAi)*, and *islet1(RNAi)* animals stained for *TPH^+^* via *in situ* hybridization 6 days after head amputation. Fused eyespots are visible after *SSDP2(RNAi)*, *LDB1(RNAi)*, or *islet1(RNAi).*Decreased peripheral *TPH^+^* puncta were noted after knockdown of *SSDP2* or *LDB1*, but not *islet1* (see insets). D) *control(RNAi)*, *islet1(RNAi)*, *SSDP2(RNAi)*and *LDB1(RNAi)* animals underwent tail amputation, 6 days of regeneration, and *in situ* hybridization with the *Wnt1* probe (Petersen and Reddien, 2008; Adell *et al.*, 2009). Only posterior ends are shown. E) *control(RNAi)*, *islet1(RNAi)*, *SSDP2(RNAi)* and *LDB1(RNAi)* animals underwent head amputation, 7 days of regeneration, and *in situ* hybridization with the *sFRP-1* probe (Gurley *et al.*, 2008; Petersen and Reddien, 2008). Only anterior ends are shown. F) *control(RNAi)*, *islet1(RNAi)*, *SSDP2(RNAi)* and *LDB1(RNAi)* animals underwent head amputation, 6 days of regeneration, and *in situ* hybridization with the *slit1* probe (Cebrià *et al.*, 2007). Animals were imaged with the dorsal side facing up; arrowheads indicate the anterior-most dorsal *slit* signal. G) Diagrams of the amputation plane used for experiments in (H). H) *control(RNAi)*, *islet1(RNAi)*, *SSDP2(RNAi)* and *LDB1(RNAi)* animals underwent sagittal amputation, 9 days of regeneration, and *in situ* hybridization with the *ChAT* probe (Nishimura *et al.*,2010). Animals are shown with the regenerating side to the right. I) We measured the area of both the newly regenerated brain lobe (right) and the existing brain lobe (left) and created a ratio for each animal in (H). *SSDP2(RNAi)* and *LDB1(RNAi)* animals regenerated significantly less new brain tissue than *control(RNAi)* animals. (n=6-11; Kruskal-Wallis with Dunn’s correction for multiple comparisons. P is an adjusted value, ** indicates P≤0.005 and **** indicates P≤0.0001). Scale=500 μm (B, C, F, H), 100 μm (B, C insets), 200 μm (D, E).

Previous research indicated that planarian *islet1* is expressed in anterior and posterior poles during regeneration (Fig. 3A, (Hayashi *et al.*, 2011; Marz *et al.*, 2013)), and we find that both *SSDP2* and *islet* are expressed broadly in regenerating tissue (Supp. Fig. 2A). Knockdown of *islet1* leads to eye fusion at the midline, failure to reestablish anterior and posterior poles, and defective gene expression at the midline (Hayashi *et al.*, 2011; Marz *et al.*, 2013). Due to the superficial similarities of the *islet1(RNAi)*, *SSDP2(RNAi)*, and *LDB1(RNAi)* phenotypes, we next investigated whether *SSDP2* and *LDB1* are required for reestablishment of polarity. *islet1(RNAi)*leads to reduced *wnt1* expression in tails after posterior amputation (Fig. 3D, (Hayashi *et al.*, 2011; Marz *et al.*, 2013)). Similarly, *SSDP2(RNAi)* or *LDB1(RNAi)* decreases *wnt1* expression in regenerating tails (Fig. 3D). We also found that *SSDP2(RNAi)* or *LDB1(RNAi)* reduces *sFRP-1* expression in regenerating heads (Fig. 3E). We conclude that SSDP2 and LDB1 likely cooperate with Islet1 in regeneration of anterior and posterior pole structures.

RNAi targeting planarian *islet-1* also leads to mediolateral patterning defects, which are reflected in regeneration of cyclopic heads and reduced expression of *slit* (Fig. 3F, (Marz *et al.*, 2013)). We confirmed that *islet1(RNAi)* specifically reduced the dorsal expression domain of slit, with a posterior shift in dorsal slit expression (Fig. 3F, arrowhead). We next investigated whether knockdown of *SSDP2* or *LDB1* phenocopied *islet1(RNAi)* impacts on *slit* expression. Indeed,*SSDP2(RNAi)* or *LDB1(RNAi)* led to reduced dorsal *slit* expression domains (Fig. 3F, arrowheads), though ventral *slit* expression was less affected. Based on these results, we expected that *islet1*, *SSDP2*, and *LDB1* might be important for lateral regeneration as well as anterior regeneration. We performed RNAi targeting *islet1*, *SSDP2*, or *LDB1* and completed sagittal amputations along the midline of the planarian body (Fig. 3G). We killed and fixed animals 9 days post-amputation and completed ISH with our *ChAT* riboprobe to investigate regeneration of internal structures (Fig. 3H). After lateral regeneration, *SSDP2(RNAi)* or *LDB1(RNAi)* led to diminished lateral regeneration, significantly smaller regenerated brain lobes, and frequently undetectable regenerated ventral nerve cords (Fig. 3H-I, Supp. Fig. 2B-C). We detected a small decrease in brain size after *islet1(RNAi)* that was not significant (Fig. 3H-I). Taken together, we conclude that planarian LDB1 and SSDP2 play roles in midline reestablishment and lateral regeneration. Although Islet1 is also a logical partner for the SSDP2/LDB1 transcriptional complex in the context of lateral regeneration, the severity of *LDB1(RNAi)* and *SSDP2(RNAi)* phenotypes indicate that other transcriptional complex partners might also be involved.

### Identification of additional planarian LIM-HD- and LMO-encoding genes

To identify other partners in LDB1/SSDP2 complexes, we first explored LIM-HD proteins. In addition to prior characterization of *islet1*, *arrowhead*, and *lhx1/5-1* (Hayashi *et al.*, 2011; Currie and Pearson, 2013; Marz *et al.*, 2013; Roberts-Galbraith *et al.*, 2016; Scimone *et al.*, 2020), a few additional LIM-HD-encoding proteins have been identified. A second homolog of Islet, *islet2*, is expressed in the anterior pole within the planarian head (Li *et al.*, 2019). *lhx2/9-1* (previously called *lhx2b*) is expressed in the planarian intestine and RNAi causes a mild reduction in goblet cells (Forsthoefel *et al.*, 2020). *lhx2/9-3* (previously *lhx2/9-like)* and *lhx3/4* (previously *lhx3/4-like)* were both identified as transcription factor-encoding genes expressed in planarian stem cells (Scimone *et al.*, 2014).

We searched available transcriptomes (Brandl *et al.*, 2016; Rozanski *et al.*, 2019) to reveal the full complement of LIM-HD- and LMO-encoding genes in *S. mediterranea*. We found that planarians express 13 genes that encode proteins with a pair of LIM domains and a homeodomain (i.e. LIM-HD proteins) (Supp. Table 3, supported by data in Molina and Cebrià, personal communication and forthcoming preprint). We also found that planarians express 3 genes that have LIM domains only and that also have homology to LIM-only (LMO) genes like *beadex*, *rhombotin*, and *lmo4* (Supp. Table 3, Supp. Fig. 3). We also identified 41 transcripts that encode proteins with LIM domains but that do not clearly belong within LIM-HD or LMO families (Supp. Table 3).

Next, we sought to characterize additional LIM-HD- and LMO-encoding transcripts by determining their expression patterns. Our ISH for published transcripts largely confirmed previous expression patterns, with the additional finding that *islet1* is strongly expressed in the planarian brain and is broadly expressed outside the poles (Fig. 4A). We found that two LIM-HD-encoding genes, *lhx2/9-1* (previously called *lhx2b*) and *lhx1/5-2*, are expressed in the planarian intestine (Fig. 4A). We noted one gene, *lmx1a/b-4*, expressed in the parenchyma, and we saw enriched expression of the majority of LIM-HD genes in subsets of cells in the planarian brain (Fig. 4A). We found that the three LMO-encoding genes had enriched expression in neurons and, for *lmo1/3-2*, in additional cell types (Fig. 4B). We saw broad concordance between our ISH results and results from single-cell sequencing analyses (Supp. Table 3 (Fincher *et al.*, 2018; Plass *et al.*, 2018)), though our ISH experiments also revealed region-specific expression patterns for several genes that were not significantly enriched in single-cell analyses. Taken together, the tissue-specific expression patterns of LIM-HD and LMO genes offer hints to potential functions of these transcription factors and additional LDB/SSDP complexes in planarian neurobiology and regeneration. Further, neural expression of *islet1* indicates that there could be additional roles still to be discovered for known LIM-HD transcription factors.

**Figure 4.**
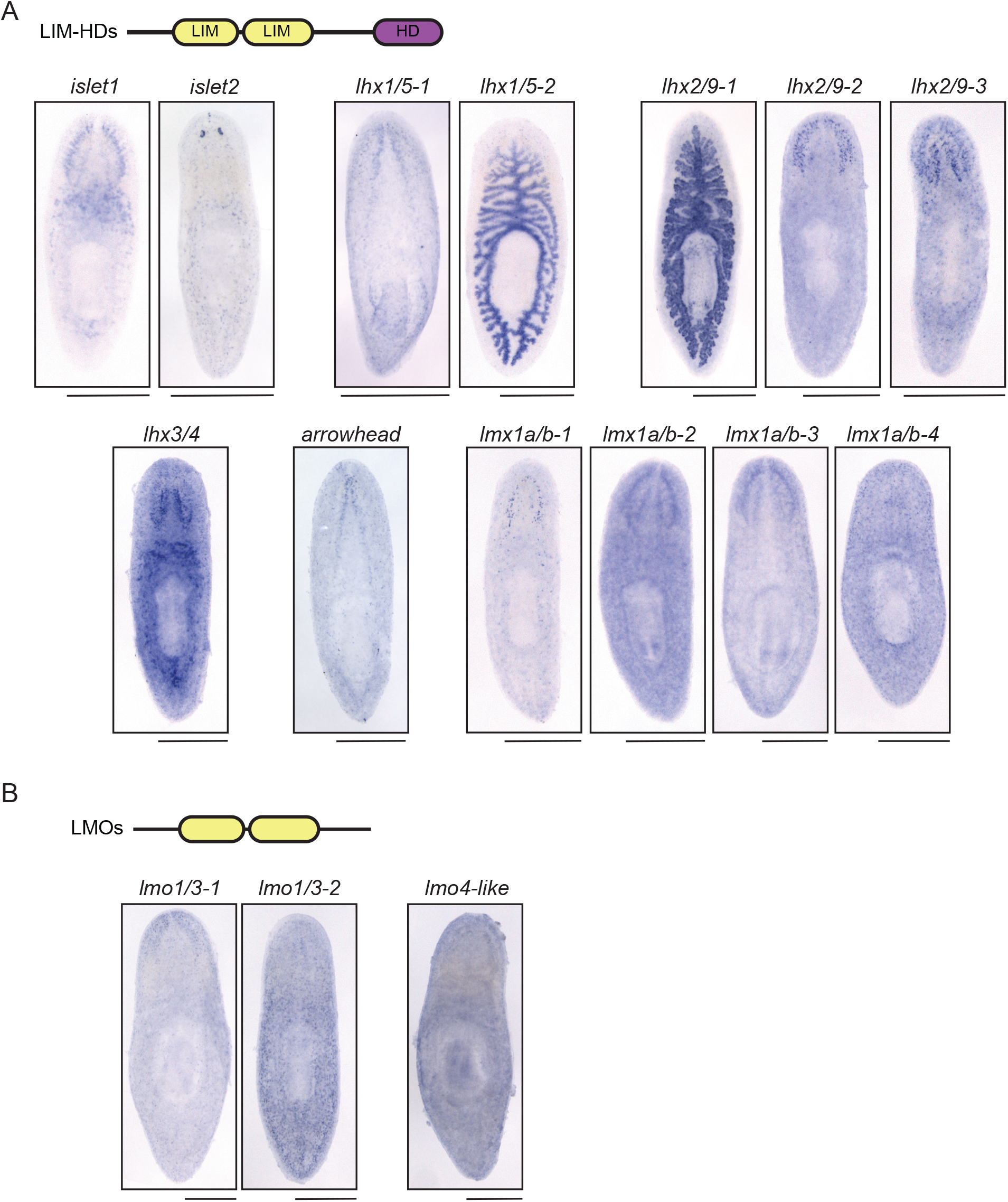
4) Expression of planarian LIM-homeodomain and LIM-only-encoding genes. A) Domain architecture of a typical LIM-homeodomain (LIM-HD) protein. 13 planarian genes encode LIM-HD proteins. Expression of these genes is shown in untreated planarians and genes are grouped by family. B) LIM-only (LMO) proteins have a typical domain architecture with two LIM domains each. 3 planarian genes encode LMO proteins. *in situ* hybridization for each LMO gene is shown. Scale=500 μm.

### Impacts of SSDP2(RNAi) on gene expression in planarians

Finally, we sought to perturb elements of the conserved LDB1/SSDP2 transcriptional complex for several reasons: 1) to uncover downstream impacts of their perturbation, 2) to discover new functions for the complex, and 3) to explore whether novel functions were accomplished through LIM-HD-dependent or independent mechanisms. We performed *SSDP2(RNAi)* in triplicate (Fig. 5A) and used bulk RNA-sequencing to identify genes differentially expressed after knockdown. We confirmed that *SSDP2* itself was 3.5x downregulated in the samples (Supp. Table 4-5). We identified 64 upregulated transcripts and 53 downregulated transcripts after *SSDP2(RNAi)* (fold change ≥ 1.5x, corrected P≤0.05; Fig. 5B, Supp. Table 4-5). A large percentage of genes downregulated after *SSDP2(RNAi)* are expressed in the intestine, in parenchymal cell types, or in *cathepsin^+^* cell types (Supp. Table 4-5, examples in Fig. 5C).

**Figure 5.**
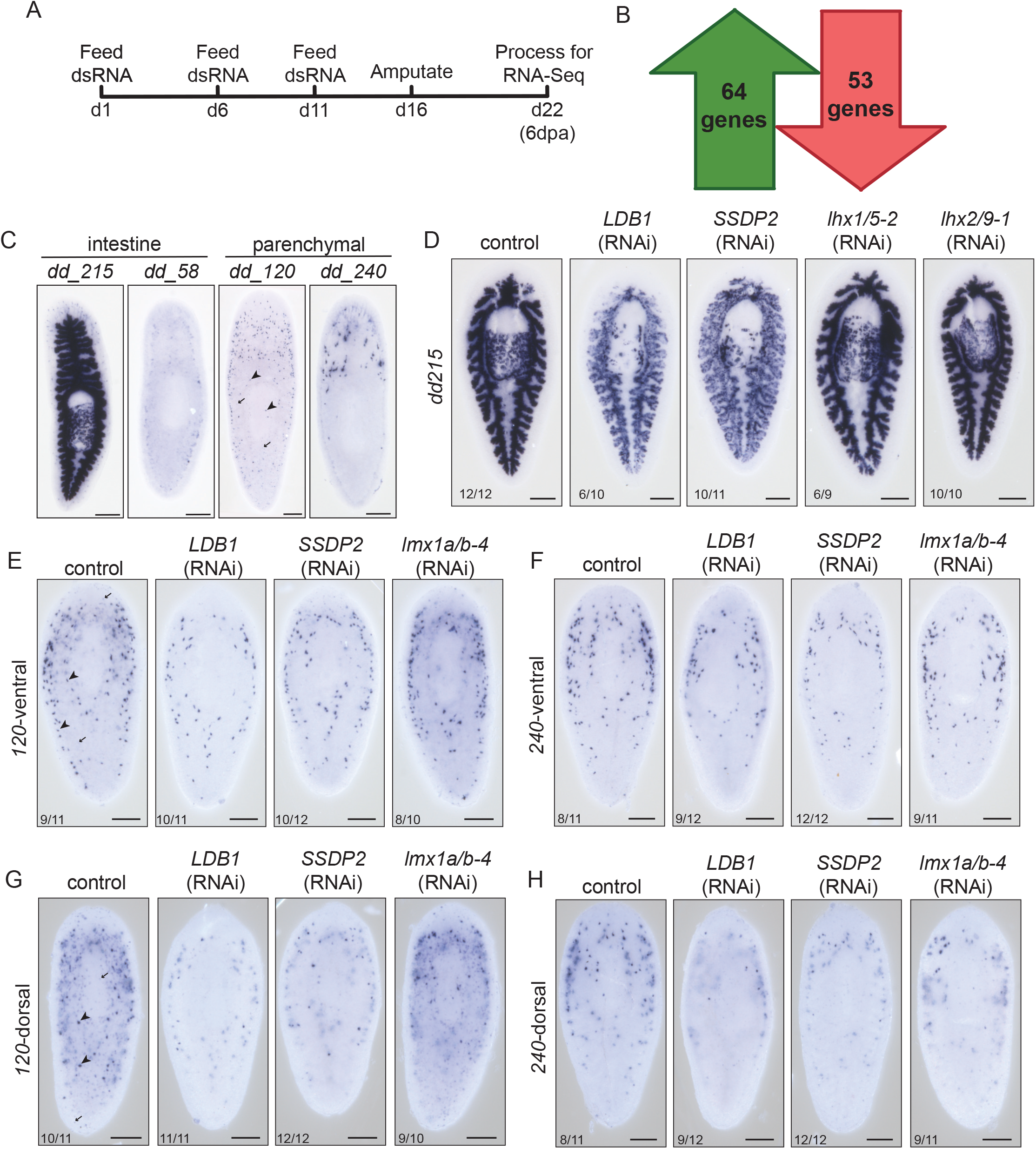
5) Identification of genes differentially expressed after *SSDP2(RNAi)*. A) RNAi paradigm used for bulk RNA-seq experiments. Animals underwent head amputation and 6 days of regeneration. B) We identified 64 genes that were upregulated ≥1.5x with a P value <0.05 after false discovery rate correction. We also identified 53 genes that were downregulated ≥1.5x with a P value <0.05 after false discovery rate correction. C) *In situ* hybridization showing normal expression of four genes significantly downregulated after *SSDP2(RNAi)*. Two parenchymal- and two intestine-enriched genes are shown. D) One transcript downregulated after *SSDP2(RNAi)* in our RNA-Seq data is *dd_215*. Using *in situ* hybridization, we show that *dd_215* is absent from the blastema and is downregulated in both the existing intestine and pharynx after either *LDB1(RNAi)*or *SSDP2(RNAi)*, but not after knockdown of the gut-enriched LIM-HD-encoding genes *lhx1/5-2* or *lhx2/9-1*. E-H) *dd_120* and *dd_240* are decreased in expression after either *LDB1(RNAi)* or *SSDP2(RNAi)*, but not after knockdown of the parenchymal-enriched *lmx1a/b-4*. Ventral (E-F) and dorsal (G-H) views of animals are shown. For *dd_120*, we noted expression in two distinct cell types with strong (arrowhead) and weak (arrow) expression. Cells with weak expression are nearly absent after *LDB1(RNAi)* or *SSDP2(RNAi)*. Animals are 6 days post-head amputation and both ventral (top) and dorsal (bottom) views are shown. Scale=200 μm.

We cloned a subset of differentially expressed genes and examined their expression after *SSDP2(RNAi)*, *LDB1(RNAi*), or RNAi targeting candidate *LIM-HD* genes. We confirmed that *prosaposin (dd_215)* is downregulated in both the intestine and the pharynx after *SSDP2(RNAi)* or *LDB1(RNAi)*, supporting roles for both complex members in *prosaposin* expression (Fig. 5D). Surprisingly, knockdown of *lhx1/5-2* or *lhx2/9-1*—the two intestine-expressed LIM-HD genes—did not alter *prosaposin* expression (Fig. 5D). We conclude that SSDP2 and LIM1 play roles in regulating intestinal gene expression, in both existing and regenerating tissue. We also examined the expression of two parenchymal-expressed genes, *dd_120* and *dd_240*, and found that their expression was decreased in *SSDP2(RNAi)* and *LDB1(RNAi)* animals (Fig. 5E-H). We hypothesized that parenchymal roles for LDB1/SSDP2 could be accomplished through the function of a parenchyma-expressed LIM-HD-encoding gene, *lmx1a/b-4.* However, neither *dd_120* nor *dd_240* exhibited altered expression after *lmx1a/b-4(RNAi)* (Fig. 5E-H).

Taken together, our work shows that SSDP2 impacts gene expression in the intestine and parenchyma, in addition to roles completed through previously described LIM-HD-encoding genes. We further find that *LDB1(RNAi)* phenocopies *SSDP2(RNAi)*impacts on intestinal and parenchymal gene expression, indicating that the two proteins likely cooperate in gene regulation in these contexts. However, we were not able to assign a LIM-HD-encoding protein partner for either intestinal or parenchymal roles, suggesting that additional LDB1/SSDP2 complex members may be important. One caveat of the RNA-Seq experiment is that we used the entire regenerating animals for our analysis; this could bias our results toward highly expressed genes as well as genes like *dd_215*, *dd_120*, and *dd_240* that showed global changes in gene expression after *SSDP2(RNAi)*, even in non-regenerating tissues.

### Conclusions and future directions

In this work, we characterized planarian homologs of LDB and SSDP and discovered that LDB1 and SSDP2 play roles in organization and polarity of new tissue during regeneration. Our results provide evidence that core transcriptional scaffolds can be targeted during regeneration to identify both new *and* conserved gene functions and to prioritize transcriptional regulators for further study. We show that LDB1 binds to SSDP2 and to LIM-HD proteins Arrowhead, Lhx1/5-1, and Islet1. We further demonstrate that LDB1 and SSDP2 are required for functions of Lhx1/5-1 and Islet1 in serotonergic neuron maturation and polarity, respectively. We identified the full complement of LIM-HD- and LMO-encoding proteins in planarians, though further work will be required to discover and characterize planarian LIM-HD proteins, LMO proteins, and other non-LIM-domain transcriptional partners of the SSDP2/LDB1 complex. Due to the expression of many LIM-HD and LMO genes in the planarian nervous system, we are particularly eager to discover roles for LDB1/SSDP2 and related factors in neuronal regeneration and function. Finally, we identify genes differentially regulated after *SSDP2(RNAi)* and discover novel roles for the transcriptional core proteins in gene expression in the intestine and parenchyma of the planarian body. We expect that future work will reveal specific DNA target sequences for LDB1/SSDP2 complexes. Taken together, our work has uncovered conserved roles for LDB/SSDP complexes in head formation and provides a starting point for further exploration of these ancient transcriptional scaffolds in animal regeneration.

## Materials and Methods

### Planarian care

We used asexually reproducing *S. mediterranea* (strain CIW4 (Newmark and Sánchez Alvarado, 2000)) for all experiments. Planarians were housed in Montjuïc salts (Cebrià and Newmark, 2005) at 18°C in the dark. We fed animals organic beef liver puree (White Oak Pastures, Bluffton, GA) approximately once per week. Animals for *in situ* hybridization or RNA interference (RNAi) were starved at least one week prior to experiments. We also used gentamicin sulfate (50 μg/mL final concentration, Gemini Bio-Products) to treat animals as needed and during all RNAi experiments.

### Sequence identification and domain analysis

We identified homologs of SSDP and LDB families using TBLASTN searches of *S. mediterranea* transcriptomes (Brandl *et al.*, 2016; Rozanski *et al.*, 2019). We used both BLAST and domain search resources at Planmine (Brandl *et al.*, 2016; Rozanski *et al.*,2019) to identify genes that encode LIM-domain proteins (LIM domain = IPR001781, PF00412). LIM-HD-encoding genes were classified as per (Molina and Cebrià, personal communication).

Translated LMO proteins were subjected to phylogenetic analysis alongside homologs of LMO proteins from *Drosophila melanogaster*, *Homo sapiens*, *Mus musculus*, *Gallus gallus*, *Danio rerio*, and *Trichoplax adhaerens* (sequences in Supp. Text 1). Human Pax1 was used as an outgroup. We completed phylogenetic analysis using the a la carte option on www.phylogeny.fr (Dereeper *et al.*, 2008). MUSCLE was used for alignment (Edgar, 2004), Gblocks was used for curation (Castresana, 2000), and PhyML was used for tree construction. 100 bootstrap replicates were performed (PhyML) and we used the WAG model of amino acid substitution (specific parameters: 4 substitution rate categories; estimates used for invariable site proportion and transition ratios).

### Molecular biology methods

For *in situ* hybridization and RNA interference experiments, we cloned 500-800 bp fragments of each gene into pJC53.2 (Collins *et al.*, 2010). Briefly, cDNA was prepared from planarians using an iSCRIPT kit (Bio-Rad) and primers shown in Supp. Table 6 were used to amplify fragments of each gene using Platinum Taq (Invitrogen). PCR amplicons were ligated into pJC53.2, which had been digested with Eam1105I, using standard ligation methods. Kanamycin was used to select transformants and sequencing was used to verify insert sequences and orientation (Azenta).

For yeast two-hybrid analysis, full length and short fragments of *LDB1*, *SSDP2*, *islet1*, *lhx1/5-1*, and *arrowhead* were amplified from cDNA with Phusion polymerase (New England Biosciences). Primers were designed to include restriction sites and are described in Supplemental Table 6. Amplicons were digested with restriction enzymes as follows: BamHI and PstI (*LDB1*, *arrowhead*, *islet1*); EcoRI/PstI (*SSDP2*); or SmaI and BamHI (*lhx1/5-1*). Digested fragments were inserted into digested pGAD424 and pGBT9 through standard molecular methods. Inserts were sequenced using primers described in Supp. Table 7.

To generate dsRNA and riboprobes, we amplified pJC53.2 inserts using a T7 primer (sequence in Supp. Table 7). PCR products were purified with a DNA clean and concentrator kit (Zymo). We synthesized dsRNA as previously described (Chong *et al.*,2013; Rouhana *et al.*, 2013). Riboprobes were generated from PCR products at 30°C overnight using either Sp6 or T3 polymerase, depending on the orientation of the insert. We included digoxigenin-11-UTP (Roche) during riboprobe synthesis. Riboprobes were treated with DNase (Promega) and purified by ammonium acetate precipitation. Riboprobes and dsRNA concentrations were determined through band intensity after gel electrophoresis, which we have found to be more reliable than a spectrophotometer as only full-length products are quantified.

### Yeast two-hybrid analysis

*Saccharomyces cerevisiae* strain PJ69-4A was used for yeast two-hybrid analysis as previously described (James *et al.*, 1996). Transformants were selected on agar plates made from dropout media (-Trp, -Leu). Interactions were evaluated on agar that was -His or -His -Ade (Clontech, now Takara Bio). Growth in the absence of either supplement indicates transcription from reconstituted GAL4 proteins. Empty vectors were used as negative controls. A pair of plasmids producing *Schizosaccharomyces pombe* Mob1 and Sid2 was used as a positive control (Salimova *et al.*, 2000; Hou *et al.*,2004).

### In situ hybridization

Whole and regenerating planarians were processed for *in situ* hybridization as described in (King and Newmark, 2013). Riboprobes produced as described above were detected using alkaline-phosphatase-conjugated anti-digoxigenin Fab fragments (1:2000, [Roche]). Colorimetric signals were generated using a development solution containing 5-Bromo-4-chloro-3-indolyl phosphate (BCIP, [Roche]) and nitro blue tetrazolium chloride (NBT, [Roche]).

### RNA interference

2-5 μg dsRNA generated as described above was combined with 30 μL of 4:1 liver:salts mix for each feeding. RNA feeding paradigms consisted of 3 feedings per experiment, with the first feeding on day 1, the second feeding on day 6, and the third feeding between days 10-12. Amputation occurred 5-7 days after the final feeding along prepharyngeal (anterior to the pharynx), postpharyngeal (posterior to the pharynx), or sagittal (down the midline) amputation planes. Animals were fixed and killed (or processed for RNA purification) 6 days post-amputation, with the exception being 9 days post-amputation for lateral regeneration experiments (Fig. 3G-H). *dsRNA* matching *Aequorea green fluorescent protein* (*GFP*) was used as a negative control in RNAi experiments.

### Microscopy

Animals that had been subjected to *in situ* hybridization were mounted in 80% glycerol or 1xPBS prior to imaging. Stained and live animals were imaged with either an Axiocam color camera mounted on a Zeiss Axio Zoom.V16 microscope or with a Leica DFC420 camera mounted on a Leica M205A stereomicroscope. Images of yeast strains (Supp. Fig. 1) were captured on an Apple iPhone 5.

### RNA sequencing and analysis

For each control and *SSDP2(RNAi)* samples, 4 plates of 10 worms were fed dsRNA as per the paradigm above. Animals were amputated prepharyngeally 5 days after the final feeding. 6 days post-amputation, animals for each sample were transferred to a 1.5 mL tube and salts were replaced with 250 μL TRIzol reagent (Invitrogen) before freezing on dry ice for storage at −80°C. RNAs were purified from each sample as per the manufacturer’s protocol, were DNAse treated (Promega), and were further cleaned with an RNA clean and concentrator kit (Zymo Research). RNAs were eluted 2x in 8 μL RNAse-free water. Three samples for each RNAi condition were submitted for quality analysis and library preparation with Illumina’s TruSeq Stranded mRNAseq kit. Libraries were sequenced on a HiSeq 4000 platform (100 nt sequencing, using sequencing kit v.1). Fastq files were demultiplexed and adaptors were trimmed. We received between 37 million and 45 million reads per sample. RNA-seq reads have been deposited as BioProject PRJNA907760 (Accession: SRX18465766-SRX18465771).

Sequences were imported into CLC Genomics Workbench 8 (QIAgen) and were mapped to the dd_Smed_v6 transcriptome (Brandl *et al.*, 2016; Rozanski *et al.*, 2019) using “batch” mode and standard parameters. We then completed gene expression comparison (EDGE) and statistical analysis, with a filter cutoff of 3 counts and with FDR correction. We present data in Supp. Table 4-5 with common and tagwise dispersion. For further analysis, we prioritized genes with P<0.05 after FDR correction and with the highest fold changes.

## Abbreviations

CLIM: co-factor of LIM domains
DNA: deoxyribonucleic acid
dpa: days post-amputation
ISH: *in situ* hybridization
LDB: LIM domain binding protein
LIM: LIN-11, Isl-1, and Mec3
LIM-HD: LIM homeodomain
LHX: LIM homeobox
LMO: LIM only
RNA: ribonucleic acid
SSDP: single-stranded DNA-binding protein

## Acknowledgements

The authors thank Dr. Phillip Newmark and past members of the Newmark laboratory for their feedback on this project in its earliest stages. We thank current and past members of the Roberts-Galbraith laboratory for feedback and support during this project. We are grateful to Drs. M. Delores (Loli) Molina Jimenez and Francesc Cebrià for helpful discussion and for sharing their work on planarian LIM-HD genes. We are grateful to Dr. Alejandro Sánchez Alvarado and Shane Merryman, of the Stowers Institute for Medical Research, for providing *S. mediterranea* during our laboratory startup and for sharing advice on planarian aquaculture. We thank Dr. Alvaro Hernandez and the staff of the Roy J. Carver Biotechnology Center (University of Illinois, Urbana-Champaign) for sequencing and experimental design advice for our RNA-sequencing experiments. We thank Dr. Kathleen Gould (Vanderbilt University) for sharing yeast strains and plasmids used in yeast two-hybrid analysis. We also thank the farmers and staff of White Oak Pastures for raising healthy cattle and providing beef liver we use to feed our planarians (Bluffton, GA). This work was supported by funding from the McKnight Foundation (McKnight Scholars Award to RRG) and the National Institute for Neurological Disease and Stroke at the National Institutes of Health (1R01NS128096-01). KBC was supported by the National Institute of General Medical Sciences of the National Institute of Health under award number 1T32GM142623 and the National Institute of Neurological Disorders and Stroke of the National Institute of Health under project number 5R25NS107179-02. KBC was also supported by the University of Georgia Research Foundation and the ARCS Foundation.

## Supplemental Figure Legends

**Supplemental Figure 1.**
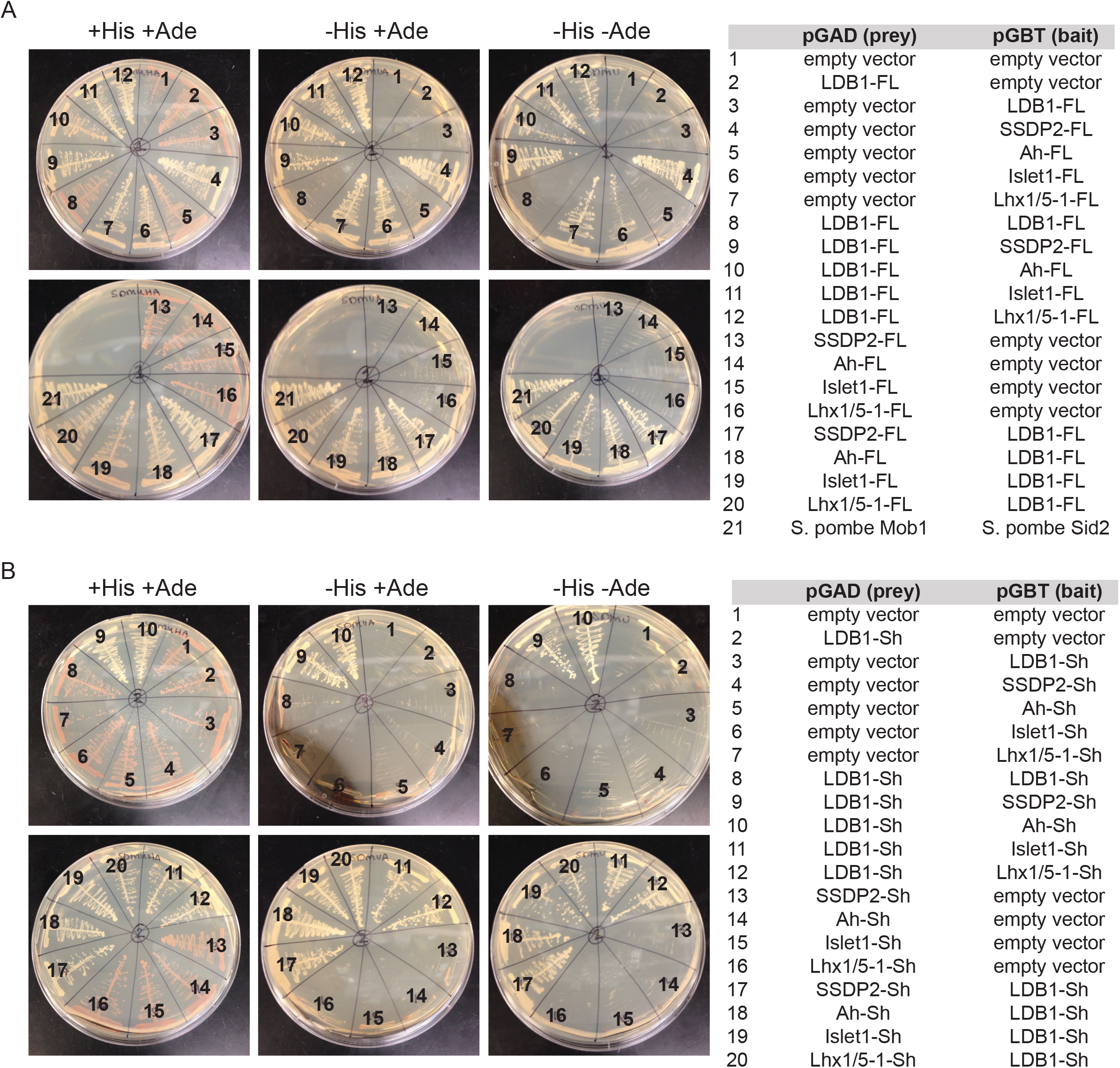
1) Yeast two-hybrid analysis of interactions between LDB1, SSDP2, and LIM-HD proteins. A-B) Strains are shown grown on selective media as indicated. Pairs of plasmids are numbered and shown to the right. (A) shows assays performed with full length (FL) proteins, while (B) indicates assays performed with shorter (Sh) constructs. See Fig. 2B-C for the summary of these data.

**Supplemental Figure 2.**
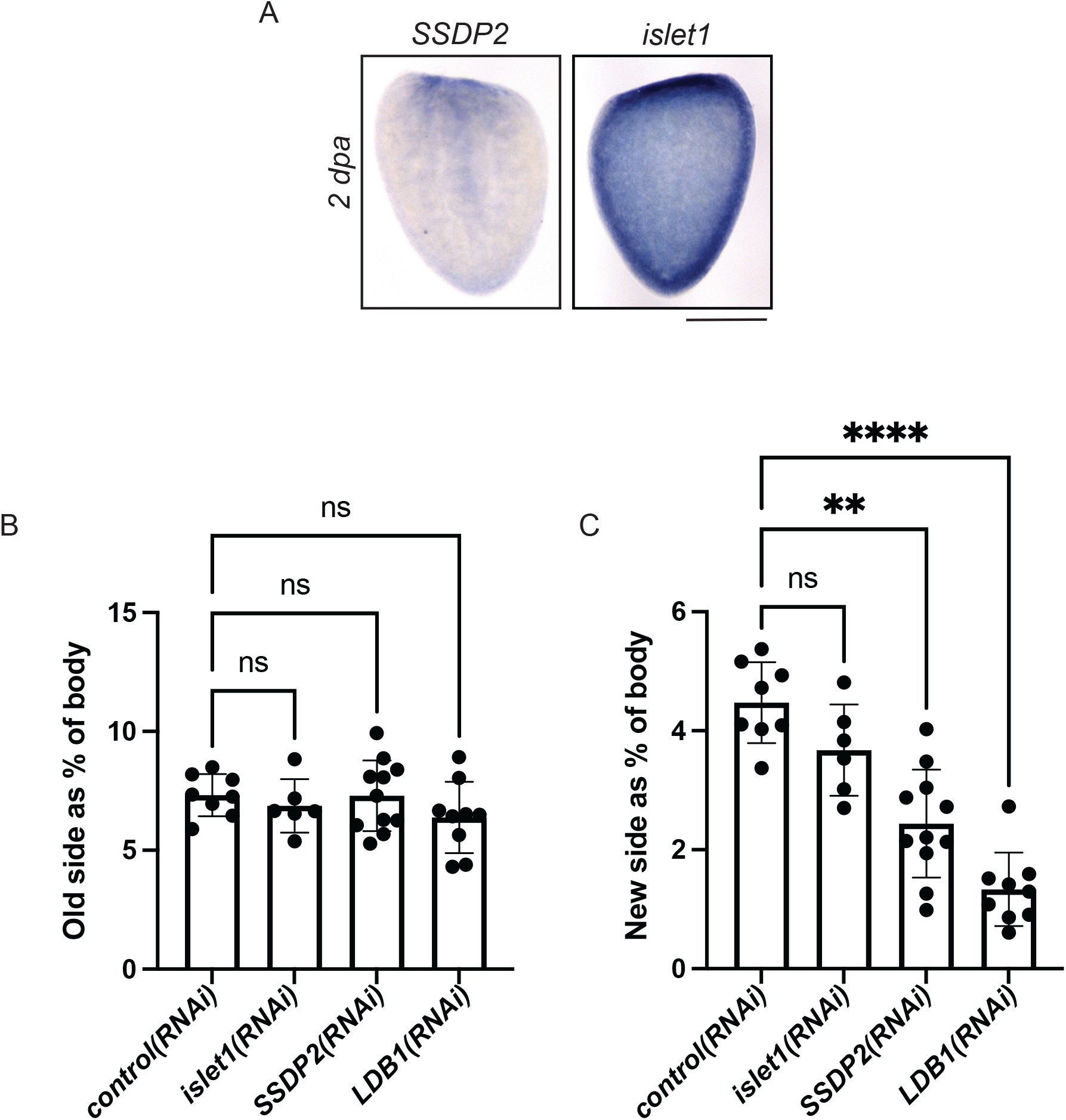
2) Additional characterization of *SSDP2* in regeneration. A) Expression patterns of SSDP2 and islet1 are shown 2 days post-amputation. B-C) For each animal from the experiment shown in Fig. 3H, we measured the area of each newly regenerated brain lobe, each existing brain lobe, and the area of the animal’s body. We created ratios for each animal in 3H to describe the old brain lobe as a ratio of the body area (Supp. 2B) and each new brain lobe as a ratio of the body area (Supp. 2C). We saw no changes in the size of the old brain lobe among any treatment group. *SSDP2(RNAi)* and *LDB1(RNAi)* animals regenerated significantly less new brain tissue than *control(RNAi)* animals, but existing brain lobes were not significantly smaller. (n=6-11; Kruskal-Wallis with Dunn’s correction for multiple comparisons. P is an adjusted value, ** indicates P≤0.005 and **** indicates P≤0.0001). Scale bar= 500 μm.

**Supplemental Figure 3.**
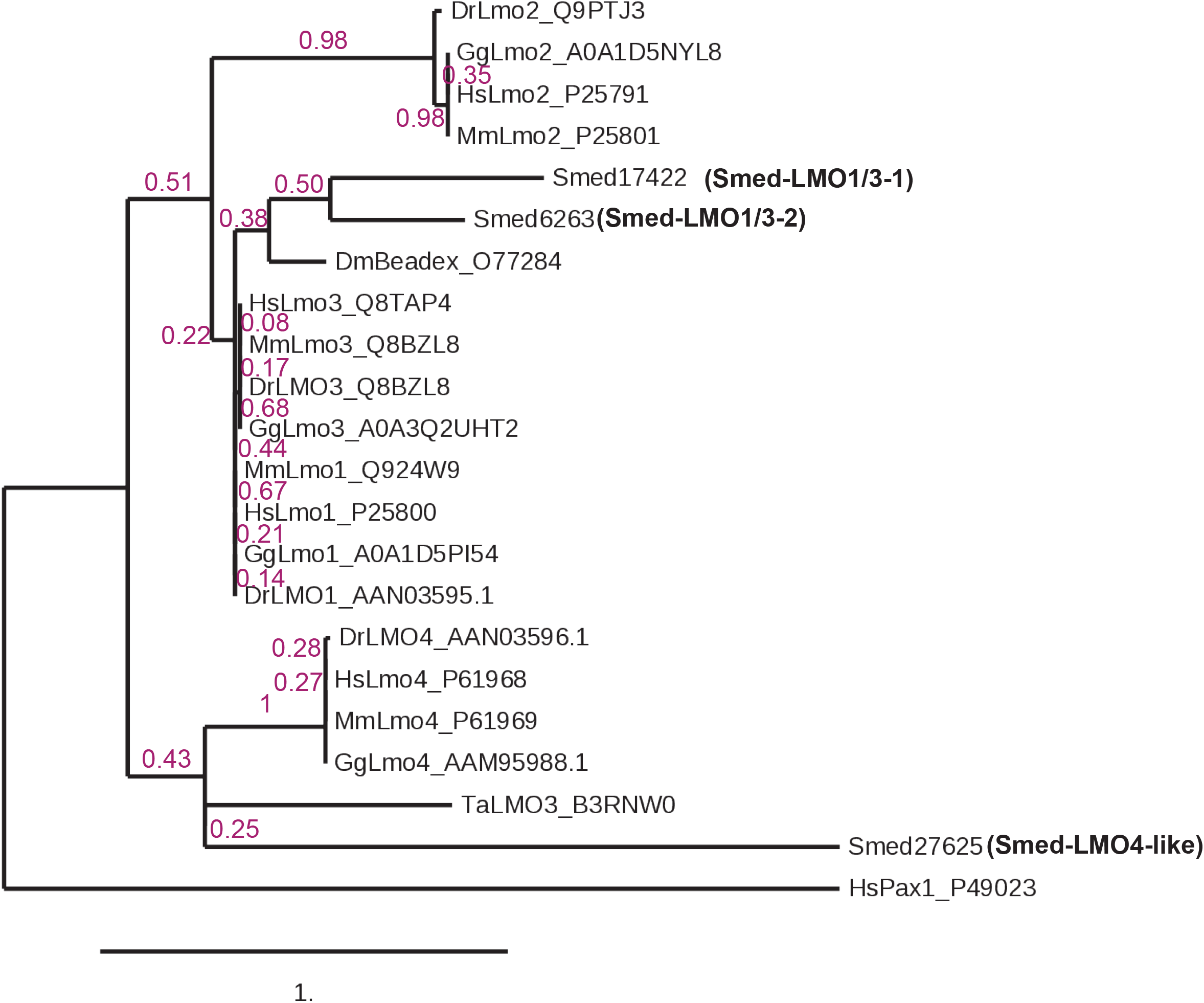
3) Phylogeny of planarian LMO proteins. A phylogram was created to depict relationships between planarian LMO proteins and LMO proteins from humans (Hs), mice (Mm), fruit flies (Dm), zebrafish (Dr), chicken (Gg), and Trichoplax (Ta). Branch support values are shown in dark magenta.

## Supplemental Tables

**1) Planarian SSDP homologs.** Information including domain organization, homologs, and cloning primers is included.

**2) Planarian LDB homologs.** Information including domain organization, homologs, and cloning primers is included.

**3) Planarian genes that encode LIM-domain proteins, including LIM-HD and LMO homologs.** Information including domain organization, homologs, and cloning primers is included.

**4) Genes significantly up- or down-regulated after *SSDP2(RNAi)*.** RNA-sequencing results (significant only) from *SSDP2(RNAi)* experiments are shown, with downregulated transcripts indicated in orange and upregulated transcripts indicated in green. *SSDP2* itself is indicated in red.

**5) Full RNA-sequencing results after *SSDP2(RNAi)*.**

**6) Cloning primer list.**

**7) Other primer list.**Includes sequencing primers and primers used for amplifying inserts from pJC53.2.

## Supplemental Text

Supplemental text contains translated protein sequences from LMO homologs as noted, with protein IDs included.

